# Community-promoted antibiotic resistance genes show increased dissemination among pathogens

**DOI:** 10.1101/2025.05.12.653433

**Authors:** David Lund, Anna Johnning, Michaela Holmström, Laleh Varghaei, Juan Salvador Inda-Díaz, Johan Bengtsson-Palme, Erik Kristiansson

## Abstract

Antibiotic resistance is increasing among bacterial pathogens, posing one of the most severe threats to future public health. A major contributor to the increasing resistance is the dissemination of mobile antibiotic resistance genes (ARGs) among bacterial communities. These genes are ubiquitously present in various environments and are especially diverse in the human gut and wastewater. Despite this, the clinical implications of the prevalence of ARGs in these bacterial communities remain unclear. In this study, we aimed to investigate how the prevalence of ARGs in human gut and wastewater microbiomes reflects their dissemination among important bacterial pathogens. To do this, we estimated the prevalence of >30,000 ARGs, including both well-known (established) and computationally predicted (latent) genes, in >6,000 metagenomic samples. From their prevalence in the human gut and wastewater, we identified four categories of ARGs: co-promoted, human gut (HG)-promoted, wastewater (WW)- promoted, and non-promoted. Our results showed that co-promoted ARGs were by far the most promiscuous, being more frequently found across multiple bacterial phyla, and more often co-localized with broad host range conjugative elements. Co-promoted ARGs were also found to be overrepresented among genes identified in multiple pathogenic species and exhibited an overall higher genetic compatibility with both pathogens and other typical residents of the human gut and wastewater microbiomes. Taken together, our results highlight the link between the promotion of ARGs in the human gut and wastewater microbiomes and their presence in human pathogens, and, thereby, the genes’ potential risk to human health.

## Introduction

The increasing antibiotic resistance among bacterial pathogens threatens human health, causing over one million deaths yearly worldwide [1]. Bacteria develop resistance as a result of changes in their genetic makeup, often through the acquisition of mobile antibiotic resistance genes (ARGs) via horizontal gene transfer [2]. Many ARGs are located on mobile genetic elements, such as transposons and plasmids, which can make them transferable between distantly related bacteria. This facilitates efficient dissemination within and between microbial communities [3, 4]. To date, thousands of ARGs that collectively confer resistance to almost every class of clinically useful antibiotics have been described and recorded in databases such as CARD and ResFinder [5, 6]. The number of known ARGs is constantly increasing as new, often more efficient, variants are discovered, typically after they have been acquired by a pathogenic host and caused hard-to-treat infections.

Bacterial communities are known to maintain a diverse array of ARGs, collectively referred to as the resistome. Investigations of the resistome utilizing shotgun metagenomics – a method wherein DNA fragments are randomly sequenced from the genomes of all microorganisms present in a sample – have demonstrated that different ARGs have complex abundance patterns that vary between environmental contexts [7-9]. Both the human gastrointestinal tract and sewage treatment plants have been demonstrated to have particularly comprehensive resistomes, encompassing ARGs that are also frequently carried by pathogenic bacteria [10, 11]. These environments create opportunities for exchange of genetic information between bacteria, potentially under selection pressure from antibiotics and other chemicals, and have therefore been suggested as potential hotspots for the promotion of ARGs.

Even though the resistome has been studied in detail for many environments, we still know little about how the composition of ARGs reflects the risks to human health. In fact, resistance genes in bacterial communities are often maintained by commensal and non-pathogenic bacteria. However, for an ARG to impact antibiotic treatments, it needs to be present and induce a resistance phenotype in a bacterium able to cause infections with adverse outcomes, especially in humans. The association of ARG abundance patterns to infection control is vital for wastewater surveillance, something that has so far been challenging [12, 13]. Recent efforts have been made to assess the risk of ARGs present in bacterial communities based on various scoring systems [14, 15]. Nonetheless, these methodologies are inherently constrained due to the absence of empirical data associating gene promotion with implications for human health. This limitation hinders risk assessment and complicates data interpretation and decision-making processes [16].

In this study, we aim to explore and clarify the connection between the prevalence of ARGs in human gut and wastewater bacterial communities and their potential risks to human health. To this end, we estimated the prevalence of >30,000 well-characterized (established) and computationally predicted (latent) ARGs in >6,000 shotgun metagenomes and >1.6 million bacterial genomes. Our results show that genes promoted in both human gut and wastewater microbiomes have a substantially higher potential to spread among human pathogens. We, furthermore, demonstrate that this potential is facilitated by the frequent presence of broad host range conjugative elements in the genetic context of these ARGs and an overall genetic compatibility between the ARGs and a large variety of bacterial hosts. Our results, thus, provide new insights into the link between the promotion of ARGs in bacterial communities and their dissemination among pathogens.

## Results

### Identification of promoted resistance genes in microbial communities

We detected the presence of 30,654 ARGs in a total of 5,630 human gut and 1,034 wastewater metagenomic samples. The ARGs included both established (i.e., well-known ARGs defined by matches to the ResFinder database) and latent variants (i.e., previously uncharacterized ARGs predicted computationally) [7] conferring resistance to aminoglycoside, beta-lactam, macrolide, fluoroquinolone, and tetracycline antibiotics. The prevalence of each ARG was estimated as the proportion of samples from the two different environments where the ARG was present (≥3 mapped reads in rarefied data, see Methods). This resulted in a total of 1,592 detected ARGs, which we classified into four categories (Fig. 1): 44 *co-promoted* ARGs that were present in ≥5% of human gut samples and ≥5% of wastewater samples; 112 *human gut (HG)-promoted* ARGs that were present in ≥5% of human gut samples but <5% of wastewater samples; 48 *wastewater (WW)-promoted* ARGs that were present in ≥5% of wastewater samples but <5% of human gut samples; and 1,388 *non-promoted* ARGs that were present in <5% of human gut samples and <5% of wastewater samples (but not completely absent from all metagenomes). Notably, all HG-promoted ARGs were also detected (≥1 read) in wastewater samples, albeit at lower abundances, whereas only 25% of WW-promoted ARGs were detected in human gut samples, though some at high abundances (Supplementary Fig. 1).

**Fig. 1.**
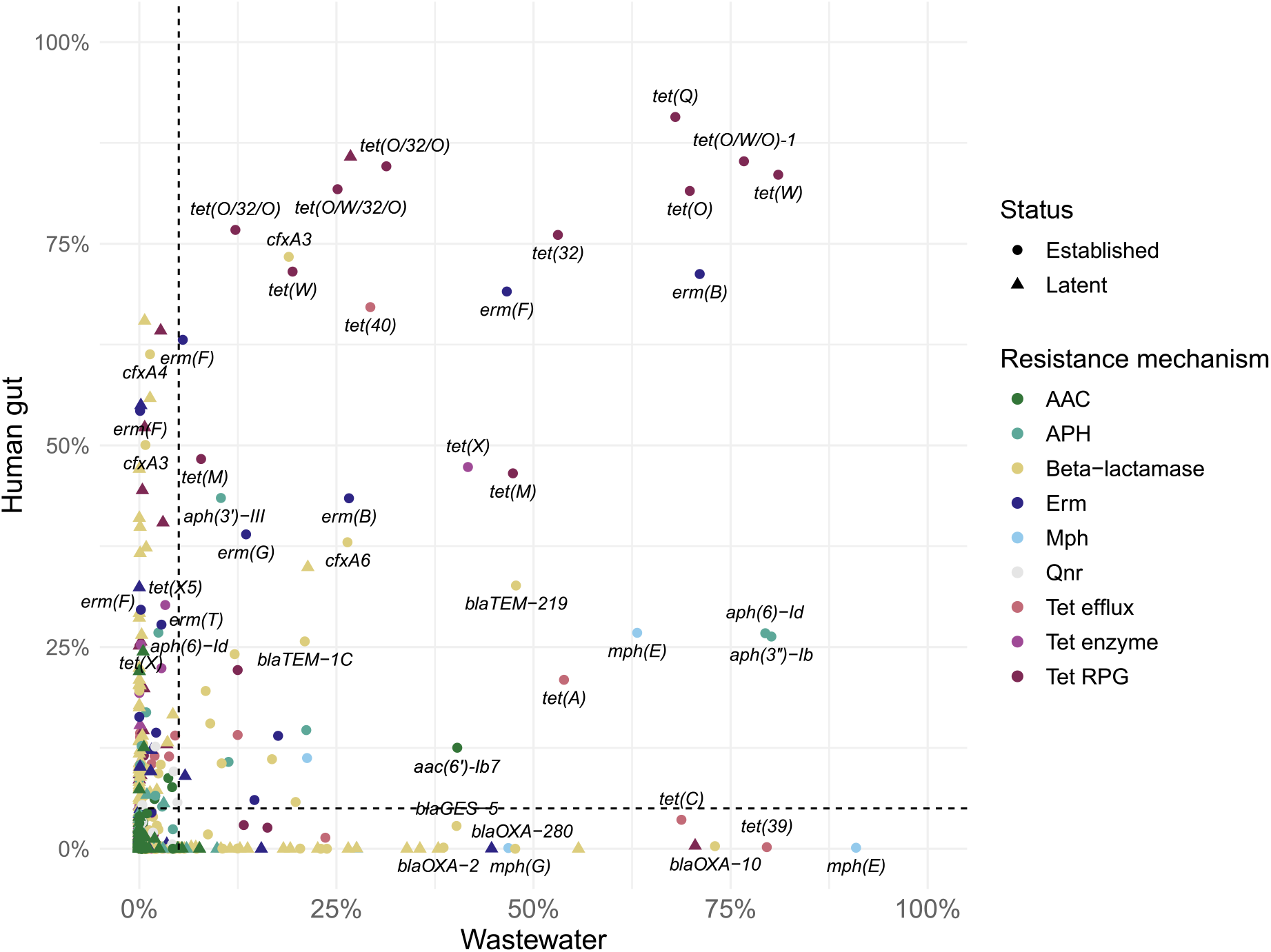
Prevalence of antibiotic resistance genes (ARGs) in human gut and wastewater metagenomic samples. Labels are included for established ARGs that were present in ≥25% of samples from either environment. The dashed lines show the classification of the ARGs into four groups: co-promoted (top right), human gut-promoted (top left), wastewater-promoted (bottom right), and non-promoted (bottom left).

Co-promoted and HG-promoted established ARGs together encompassed the largest diversity of resistance mechanisms (Fig. 2a). In particular, tetracycline resistance was significantly overrepresented among both co-promoted (RPG, *p*=3.31×10^−9^, Fisher’s exact test) and HG-promoted established ARGs (inactivating enzymes, *p*=1.87×10^−5^) Fisher’s exact test, Supplementary Fig. 2) while beta-lactamases were underrepresented (*p*=2.31×10^−3^ and *p*=1.00×10^−4^ for co-promoted and HG-promoted established ARGs, respectively). Co-promoted ARGs also included aminoglycoside (*aac* and *aph*) and macrolide resistance genes (*erm* and *mph*), but not any *qnr* genes, which were only found among the HG-promoted (established) and non-promoted ARGs (both latent and established).

**Fig. 2.**
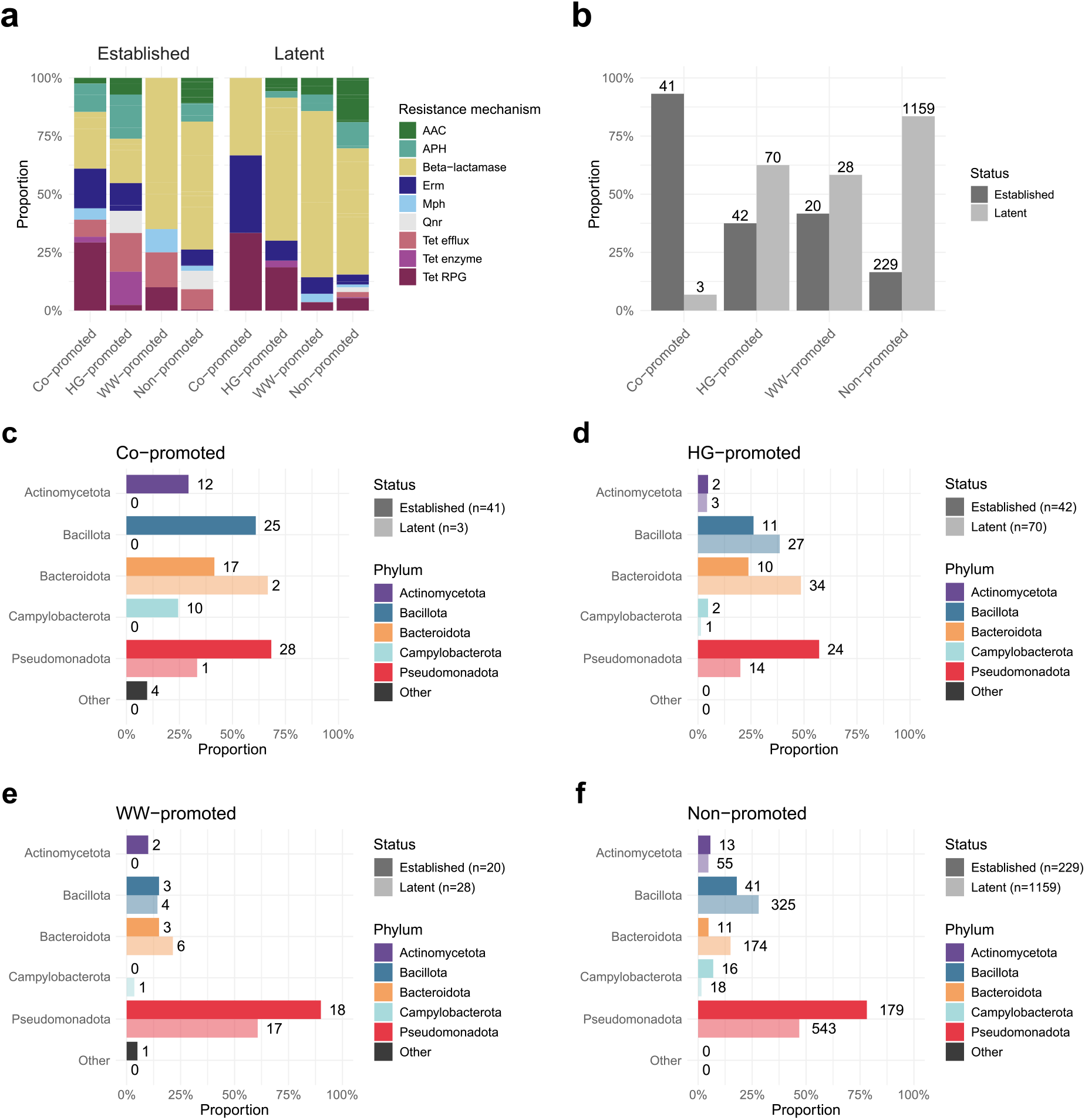
Overview of the four categories of antibiotic resistance genes (ARGs): Co-promoted, human gut (HG)-promoted, wastewater (WW)-promoted, and non-promoted. **a** Distribution of gene classes among the different categories of established and latent ARGs. **b** Proportion of established and latent ARGs comprising the different categories. Absolute numbers are displayed above each bar. **c**–**f** Distribution of phyla among the bacterial hosts carrying co-promoted, HG-promoted, WW-promoted, and non-promoted ARGs, respectively, divided between established and latent ARGs. The length of each bar represents the proportion of ARGs from that category that were identified in at least one genome from that phylum. Absolute numbers are displayed with each bar.

The co-promoted ARGs almost exclusively included established genes known to be mobile and widespread (Fig. 1, Fig. 2b). This included, for example, *tet(Q)* and *erm(B)*, as well as *aph(3’’)-Ib* (*strA)* and *aph(6)-Id* (*strB*), which are frequently found on mobile genetic elements [17]. By contrast, the HG-promoted and WW-promoted groups consisted of a more equal mix of established and latent ARGs (Fig. 2b). Established HG-promoted ARGs included e.g. *erm(F)* and *cfxA* beta-lactamases, while established WW-promoted ARGs included e.g. *tet(C)* and OXA beta-lactamases (Fig. 1). Non-promoted ARGs were highly dominated by latent resistance genes (Fig. 2b), but also included established ARGs from 21 of the 22 analyzed gene classes (all but tetracycline inactivating enzymes).

### Co-promoted resistance genes have a higher potential for horizontal transfer

Resistance genes from all four categories, both latent and established, were most frequently detected in the phyla Pseudomonadota, Bacillota, and Bacteroidota (Fig. 2c–f). However, the hosts of co-promoted and HG-promoted ARGs showed a higher diversity with a more even distribution between the major bacterial phyla. In contrast, hosts carrying WW-promoted and non-promoted ARGs primarily belonged to Pseudomonadota.

To further assess the mobility potential for ARGs promoted in the different environments, we retrieved the taxonomic affiliations of the bacterial genomes in which they were identified and obtained their hosts’ lowest common ancestor (Fig. 3). Interestingly, co-promoted genes were significantly overrepresented among established ARGs identified in multiple phyla (*p*=0.002). Indeed, as many as 49% of the co-promoted established ARGs were identified in more than one phylum, compared to 17%, 20%, and 11% of HG-promoted, WW-promoted, and non-promoted established ARGs, respectively. In contrast, HG-promoted and non-promoted established ARGs were mainly restricted to the genus level, suggesting either that these genes are relatively immobile or that they have a narrow host range (Fig. 3a). However, WW-promoted ARGs, which were largely confined to the Pseudomonadota phylum, were significantly overrepresented among all ARGs found in multiple classes within the same phylum (*p*=0.004, Fig. 3a–b). Thus, while co-promoted and HG-promoted established ARGs were observed in taxonomically similar bacterial hosts, co-promoted and WW-promoted established ARGs showed similarities in their ability to spread over large phylogenetic distances. Among latent genes, however, inter-phyla transfers were much rarer. Indeed, only the HG-promoted category contained a notable proportion of latent ARGs observed in multiple phyla (14%), while inter-phyla transferred latent ARGs were almost completely absent from the other categories (0%, 0%, and 0.2% for co-promoted, WW-promoted, and non-promoted latent ARGs, respectively).

**Fig. 3.**
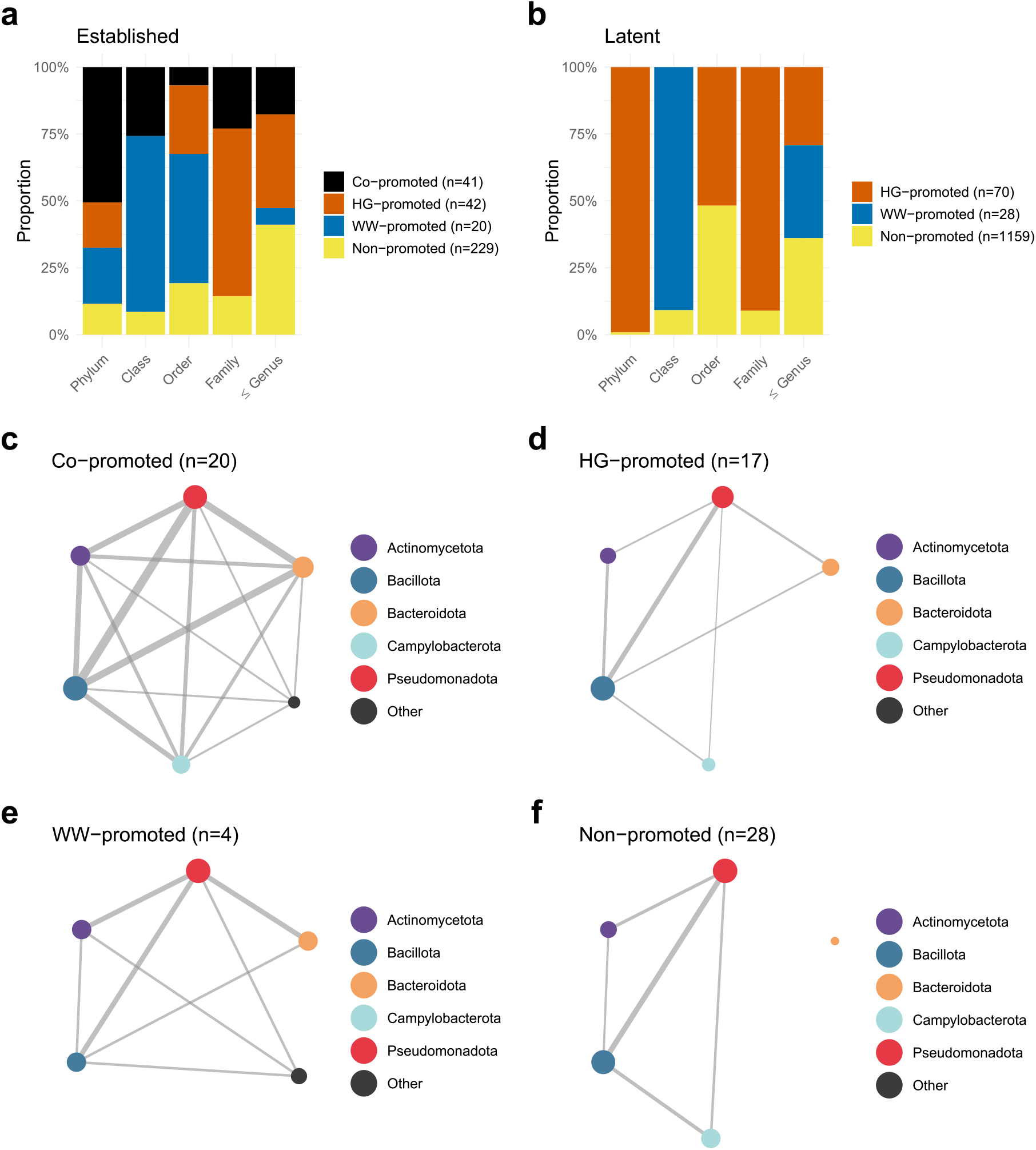
Overview of the horizontal transfer of co-promoted, human gut (HG)-promoted, wastewater (WW)-promoted, and non-promoted antibiotic resistance genes (ARGs). **a**–**b** Taxonomic host range associated with established and latent ARGs with different promotion categories. Each bar represents a taxonomic level, displaying the mean proportions of ARGs from each promotion category that transferred across no more than that taxonomic distance (based on 1,000 repetitions of subsampling the categories to equal size, see Methods for details). In panel b, co-promoted latent ARGs are excluded due to their low numbers (n=3). **c**–**f** Networks describing the transfer patterns of intra-phyla transferred ARGs with different promotion categories. Each node represents a bacterial phylum, with a size proportional to the number of inter-phyla transferred ARGs found in that phylum. Edge thickness indicates the proportion of inter-phyla transferred ARGs in a category found in both phyla (only edges representing ≥10% shown).

Network analysis was used to explore the transfer patterns for the established and latent ARGs present in multiple phyla (Fig. 3c–f). Co-promoted ARGs were found to have consistently transferred between all the major bacterial phyla, including Actinomycetota, Bacillota, Bacteroidota, Campylobacterota, and Pseudomonadota. By contrast, HG-promoted, WW-promoted, and non-promoted ARGs showed overall lower connectivity, primarily indicating transfers between Pseudomonadota and either Bacillota, Acinomycetota, or Bacteriodota (though this was rare for non-promoted ARGs). Indeed, while inter-phyla transferred co-promoted ARGs were present in an average of 4.45 phyla, other ARG categories were much more restricted (average 2.39, 3.25, and 2.39 phyla for inter-phyla transferred HG-promoted, WW-promoted, and non-promoted ARGs, respectively; Supplementary Fig. 3).

To further explore the potential for horizontal gene transfer, we analyzed the genetic contexts of ARGs in different promotion categories. The up- and downstream sequences around the co-promoted, HG-promoted, and WW-promoted ARGs were extracted from their respective host genomes and annotated for genes associated with conjugative elements, including mating pair formation (MPF) genes and relaxases, as well as co-localized mobile ARGs (Fig. 4, Supplementary Fig. 4). There was a clear difference in the diversity of MPF genes, especially between co-promoted and WW-promoted established ARGs (*p*=0.001, Fisher’s exact test, Fig. 4). This included, particularly, a higher frequency of class FATA MPF genes located close to co-promoted ARGs, while class T MPFs were more commonly located close to WW-promoted ARGs (Fig. 4). Among co-localized relaxases, MOB_P_ dominated for all promoted ARGs, however, co-promoted established ARGs were also commonly associated with MOB_V_ (Supplementary Fig. 4c). For the established ARGs, co-localization of at least one other ARG was equally frequent for the co-promoted (82.9%), WW-promoted ARGs (75%), and HG-promoted (73.8%) genes, while it was less common for non-promoted genes (28.8%, Supplementary Fig. 4e).

**Fig. 4.**
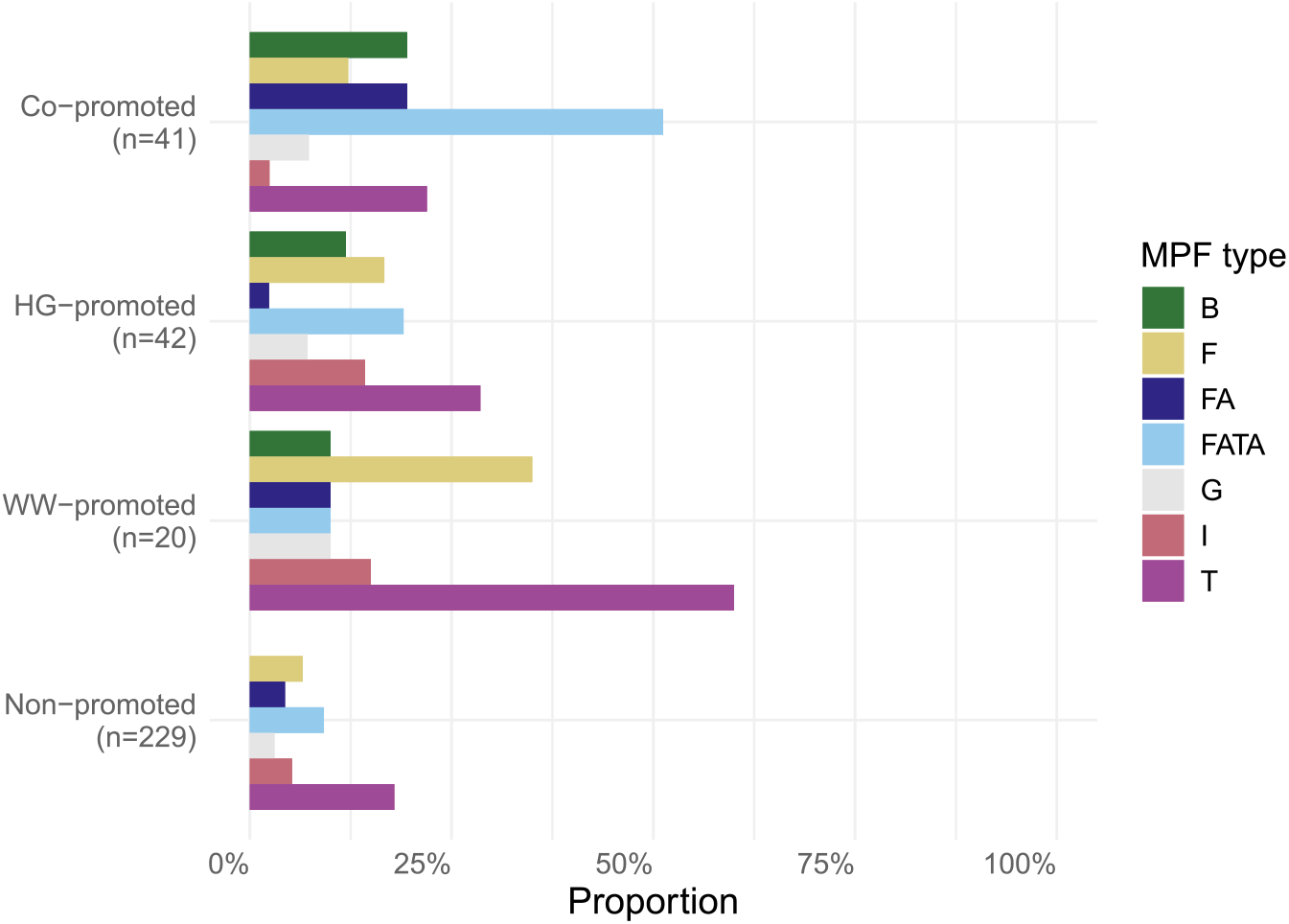
Distribution of mating pair formation (MPF) genes co-localized with established co-promoted, HG-promoted, WW-promoted, and non-promoted ARGs. For each promotion category, bar lengths indicate the proportion of ARGs with a co-localized MPF gene type detected in at least one host genome.

### Co-promoted resistance genes are more widespread in pathogens

Next, we assessed the dissemination of ARGs in different promotion categories among 15 clinically important pathogens known to be associated with high mortality [18]. We observed a strong overrepresentation of co-promoted genes among established ARGs identified in >5 pathogens (Fig. 5a, Supplementary Fig. 5). In fact, the vast majority (>90%) of the established ARGs that had spread to at least eight different pathogens were co-promoted. Accordingly, the co-promoted ARGs was the only category commonly present in pathogens from multiple phyla, including Pseudomonadota (e.g. *Escherichia coli, Klebsiella pneumoniae*, and *Salmonella enterica*), Bacillota (e.g. *Enterococcus spp*., *Staphylocoocus aureus*, and *Streptococcus spp*.), and Campylobacterota (*Campylobacter jejuni*) (Fig. 5c–f). In contrast, HG and WW-promoted established ARGs were common in pathogens from Pseudomonadota, but rare in other phyla. WW-promoted ARGs were, however, somewhat more common among established ARGs identified in ≤5 pathogens (Fig. 5a), which reflects their high frequency in a limited number of specific pathogens (Fig. 5e).

**Fig. 5.**
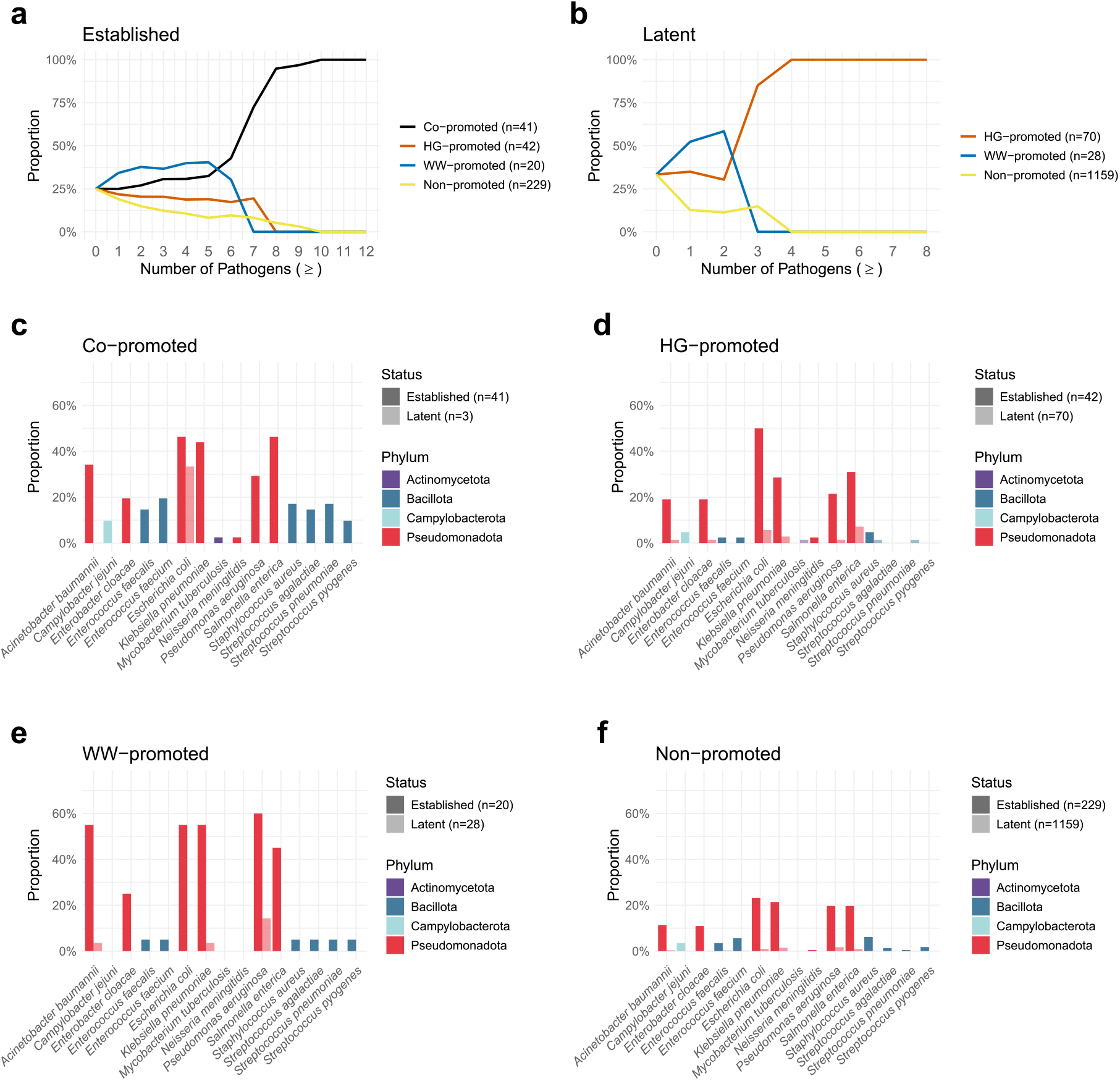
Distribution between pathogenic host species of antibiotic resistance genes (ARGs) among co-promoted, HG-promoted, WW-promoted, and non-promoted ARGs. **a**–**b** Mean proportions of established and latent ARGs carried by ≥n pathogens belonging to each promotion category (based on 1,000 repetitions of subsampling the groups to equal size, see Methods for details). In panel b, co-promoted latent ARGs are excluded due to their low numbers (n=3). **c**–**f** Proportion of ARGs from each category that were carried by each of the selected pathogens, divided between established and latent genes.

The lack of genetic compatibility has been suggested as a major barrier to the spread of ARGs between bacterial hosts [19, 20]. For each ARG and pathogenic species, we therefore calculated the difference in nucleotide composition (Euclidean distance in 5-mer distributions) between the gene sequence and a representative genome for the pathogen. Interestingly, co-promoted established ARGs had a nucleotide composition that was on average more similar to the genomes of the included pathogens, suggesting an overall higher degree of genetic compatibility (Fig. 6, Supplementary Fig. 6). The genetic dissimilarity was significantly higher for all other established gene categories when compared to the co-promoted established ARGs: WW-promoted (*p*=2.14×10^−4^, one-sided Wilcoxon’s signed rank test), HG-promoted (*p*=3.05×10^−5^), and non-promoted (*p*=1.56×10^−4^). To investigate if the differences in genetic compatibility could be observed beyond the included pathogens, we identified 110 bacterial genera that were present in ≥5% of the human gut and/or wastewater metagenomic samples and estimated their nucleotide composition dissimilarity with the differently promoted established ARGs (Supplementary Fig. 7). Our results suggested that co-promoted ARGs are more genetically similar to the residents of human gut and wastewater microbiomes. Indeed, when considering the identified resident genera, co-promoted genes showed the highest average genetic compatibility with 58% of them, in comparison with 0%, 22%, and 20% for HG-promoted, WW-promoted, and non-promoted ARGs, respectively. Finally, we observed a low but significant negative correlation between the average nucleotide composition dissimilarity of an established ARG to typical residents of the human gut and wastewater and the ARGs prevalence in the metagenomes (Spearman’s *ρ*=-0.28, *p*=1.81×10^−7^, Supplementary Fig. 8).

**Fig. 6.**
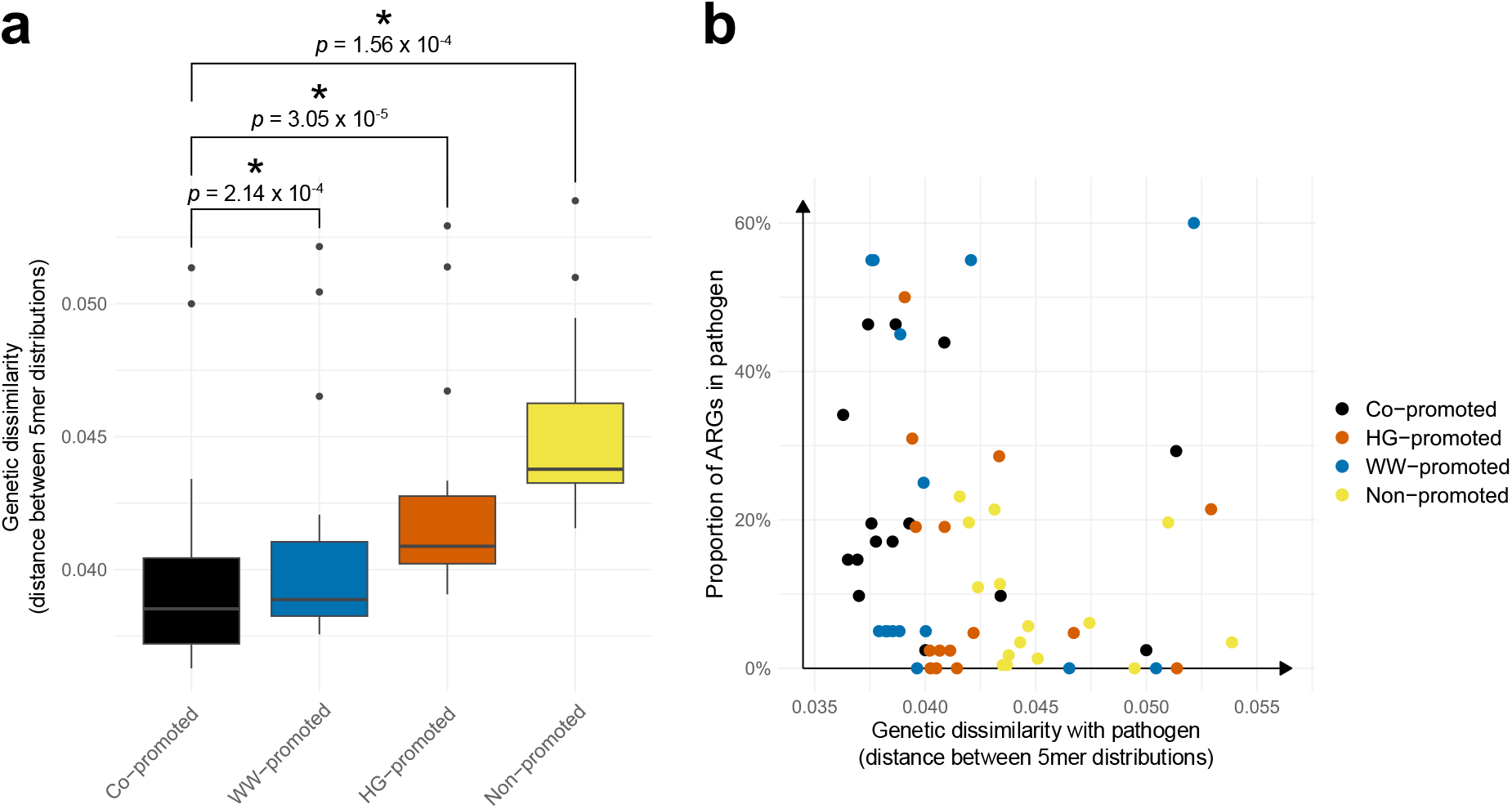
Genetic dissimilarity, estimated as the Euclidean distance between 5mer-distributions, between antibiotic resistance genes (ARGs) in different promotion categories and important bacterial pathogens. **a** Boxplots showing the distribution of median genetic dissimilarity between the ARGs from different categories and each of the included pathogens. *P*-values have been generated using Wilcoxon’s signed rank test (one-sided), comparing co-promoted ARGs to each of the other categories (\**p*<0.01). **b** Relationship between proportion of ARGs in selected pathogens and genetic dissimilarity with pathogenic host species. Each point represents one of the selected pathogens and category of established ARGs: co-promoted, human gut (HG)- promoted, wastewater (WW)-promoted, or non-promoted. The *y*-axis shows the proportion of ARGs in that category carried by that specific species, while the *x*-axis indicates the median genetic dissimilarity between the ARGs and the representative genome of the pathogen.

## Discussion

In this study, we investigated ARGs highly prevalent in the human gut and wastewater microbiomes and analyzed their potential for horizontal dissemination. Based on over 30,000 resistance genes identified in over 6,000 metagenomes and over 1.6 million bacterial genomes, we show that co-promoted ARGs, i.e., resistance genes that were re-occurring in both human gut and wastewater bacterial communities, are especially widespread among taxonomically diverse bacteria, including pathogens. In particular, co-promoted ARGs were found to co-localize with a broader diversity of conjugation genes and be more genetically compatible with both common pathogens and typical residents of the human gut and wastewater microbiomes. This study clarifies the link between ARGs promoted in human gut and wastewater microbiomes, their distribution among pathogens and, thereby, their connection to human health risks.

Our results show that co-promoted ARGs, which almost exclusively encompassed well-known (established) genes, generally have a wider host range compared to other resistance genes detected in the human gut and wastewater metagenomes (Fig. 3). In fact, as many as 49% of the co-promoted established ARGs were identified in multiple bacterial phyla, supporting at least one inter-phyla transfer event during their evolutionary history. In contrast, only 17% of HG-promoted, 20% of WW-promoted, and 11% of non-promoted established ARGs were identified in multiple phyla. Our results, thus, suggest that co-promoted ARGs have an increased potential to spread horizontally between evolutionarily divergent bacteria. Interestingly, this potential was not clearly reflected by association with plasmids and other conjugative elements. While we found that co-promoted established ARGs were co-localized with several types of conjugative systems [21], this was also the case for established resistance genes promoted solely in the human gut or wastewater bacterial communities (Fig. 4). However, co-promoted established ARGs were, to a greater extent, associated with conjugative elements carrying mating pair formation (MPF) genes from class FATA, which are almost exclusively found in bacterial hosts from Bacillota and Actinomycetota [22]. The presence of co-promoted genes in these particular phyla was further supported by taxonomic analysis of their hosts, both in general (Fig. 2c) and for pathogens (Fig. 5). In comparison, co-localization of MPF_FATA_ was rare for HG-promoted and WW-promoted established ARGs, despite the strong presence of HG-promoted ARGs in Bacillota (Fig. 2d). The HG-promoted and WW-promoted established ARGs were instead more commonly associated with MPF genes from classes T and F. Since MPF_T_ and MPF_F_ are most often identified in Pseudomondota [23], like the WW-promoted ARGs themselves, this could explain why these ARGs were less likely to transfer between phyla, yet frequently found across different classes within Pseudomonadota. Our results, thus, suggest that the higher potential to spread over large evolutionary distances observed for co-promoted ARGs is facilitated by their frequent location on specific conjugative elements that, together, provide an increased host range covering all major phyla.

Horizontally transferred genes need to be genetically compatible with their new host to confer a fitness advantage. More specifically, any acquired gene needs transcription and translation efficiency to ensure an appropriate expression level [24]. The protein must also be successfully integrated into the cellular biochemical context to induce a sufficiently strong phenotype [25]. The lack of genetic compatibility between an acquired ARG and its new host has been suggested as a significant barrier that can hinder efficient spread between genetically diverse bacteria [26]. Interestingly, our results indicate that co-promoted established ARGs generally have higher genetic compatibility with important bacterial pathogens than other resistance genes detected in the human gut and wastewater microbiome. This higher compatibility, which was estimated as the difference in nucleotide composition, varied in size but was apparent for all investigated pathogens regardless of phylum, with the exceptions of the GC-rich *Mycobacterium tuberculosis* and *Pseudomonas aeruginosa* (Fig. 6b, Supplementary Fig. 6). A similar trend could be seen for HG- and WW-promoted established ARGs, which had overall higher compatibility with pathogens than non-promoted resistance genes. The co-promoted established ARGs also showed an overall higher genetic compatibility with common residents of the human gut and wastewater microbiomes, although there was significant variability between genera (Supplementary Fig. 7). Our results, thus, suggest that the higher genetic compatibility of co-promoted ARG may contribute to their ability to transfer efficiently, especially between evolutionarily divergent bacterial hosts [19]. There were, however, also established ARGs in the analyzed environments that were rare in pathogens, despite displaying high genetic compatibility with these bacteria. This suggests that compatibility alone – at least not in the way it is captured in this study – is insufficient to allow widespread dissemination among clinically important bacterial pathogens.

The strong association between the promotion of ARGs in microbial communities and their increased potential to spread horizontally suggests that these processes are entwined. Indeed, the human gut and wastewater microbiomes both contain complex mixtures of bacteria, which allows for horizontal transfer of resistance genes between divergent hosts, sometimes under strong selection pressures [4, 27-30]. Furthermore, there is mounting evidence that these and other environmental bacterial communities act as reservoirs from which resistance genes can be recruited. This is exemplified by the many clinically relevant ARGs that have been transferred to pathogens from diverse non-pathogenic bacteria, some of which are abundant in the human gut and wastewater treatment plants [31, 32]. Our results, therefore, emphasize the need for a holistic perspective to fully understand the processes behind the mobilization, acquisition, promotion, and dissemination of antibiotic resistance genes. Interestingly, we noted that the co-promoted ARGs most widely spread among pathogenic species encompassed genes known to have circulated among pathogens for decades. These included, among others, the beta-lactamase gene *bla*_*TEM-1*_, the tetracycline ribosomal protection gene *tet(M)*, and the macrolide resistance gene *erm(B)* [33-35]. These genes all have a long history of being maintained by bacteria under selection pressure, including natural residents of the human gut and wastewater communities and/or pathogens under antibiotic treatment [36-38]. Our observations, thus, suggest that the promotion and dissemination of ARGs across environments is a slow process; indeed, these genes were all found in pathogens more than 40 years ago. It is, therefore, likely that more recently mobilized resistance genes are currently undergoing this evolutionary process, which may increase their future clinical relevance. We, thus, argue that actions aiming to reduce the dissemination of ARGs between microbial communities should be considered when designing management strategies to meet the growing threat of antibiotic resistance.

Bacteria harbor a vast diversity of resistance genes, of which only a small part is represented in current sequence repositories [7, 39]. Our results confirm that there is a wide range of uncharacterized resistance genes – here referred to as latent ARGs [7] – that are commonly detected in both the human gut and wastewater bacterial communities. Furthermore, many of the detectable latent ARGs were promoted in human or wastewater bacterial communities. Interestingly, when looking at the genetic compatibility between latent ARGs and pathogens, we again found that promoted genes were generally more compatible with these pathogens compared to non-promoted ARGs (Supplementary Fig. 6), suggesting that they are likely under similar selection pressures as established resistance genes. Several of the promoted latent ARGs were also present close to genes associated with mobile genetic elements and were found to have spread between evolutionarily divergent bacteria. These included pathogens such as *E. coli*, suggesting that some latent ARGs may constitute emerging risks for human health. It is important to emphasize that even though our analysis was based on 33,944 ARGs identified in 1,631,873 bacterial genomes – nearly a tenfold expansion of current databases [5] – it still likely only reflects a small part of the resistome. Similarly, the analysis of human gut and wastewater microbiomes was based on 6,000 metagenomic samples, thus not reflecting the total diversity of these microbial communities. Thus, this suggests that there are potentially other, not yet discovered, resistance genes promoted in human and wastewater microbiomes.

## Conclusions

This study clarifies the link between the promotion of ARGs in bacterial communities and their dissemination among pathogens. Specifically, ARGs that frequently recur in human gut and wastewater environments were shown to be especially mobile, with approximately half of these genes having undergone inter-phyla transfers at least once. Moreover, these genes were significantly overrepresented among resistance genes carried by multiple pathogens and exhibited a higher degree of genetic compatibility with both these pathogens and other common residents of the analyzed bacterial communities. Consequently, this study provides empirical evidence concerning the correlation between the prevalence of ARGs in human gut and wastewater microbial communities and the potential implications for human health. The findings will therefore enhance the interpretation of wastewater surveillance, bridge existing knowledge gaps in risk assessments, and enable more informed and effective policymaking to address the escalating threat of antibiotic-resistant bacteria.

## Methods

### Creation of the ARG reference database

A total of 2,015,254 bacterial genomes were downloaded from NCBI Assembly (2024-03-24) [40]. To ensure the robustness of the downstream analysis, 383,381 genomes that did not pass NCBI’s taxonomy check and/or where contamination was suspected, were removed from consideration. The remaining 1,631,873 genomes were screened for ARGs using fARGene v0.1 [41], a software that uses probabilistic models to identify ARGs in bacterial genomes. In addition to known resistance determinants, fARGene also predicts uncharacterized homologs with high accuracy, as previous studies have shown [42-45]. fARGene was executed using 22 profile hidden Markov models built to identify genes conferring resistance to six major classes of antibiotics: aminoglycosides (AAC aminoglycoside acetyltransferases (six models), APH aminoglycoside phosphotransferases (three models)), beta-lactams (class A, B1/B2, B3, C, and D (two models) beta-lactamases), macrolides (Erm 23S rRNA methyltransferases (two models), Mph macrolide 2’-phosphotransferaes), tetracyclines (tetracycline efflux pumps, inactivating enzymes, and ribosomal protection genes (RPGs)), and fluoroquinolones (*qnr*). For each model, previously reported threshold scores were used [42-46], and each hit above the specified threshold was recorded as a putative ARG and stored for further analysis. This resulted in 6,834,484 hits encoding 149,036 unique protein sequences.

The ARGs predicted by fARGene were complemented with 4,150 gene sequences downloaded from the CARD database (2024-06-17) [5] that represented the same resistance gene classes as the fARGene models. To remove ARGs whose open reading frame overlapped with insertion sequences (ISs), we used BLASTx v2.15.0 [47] to align the resistance genes against 8,417 ISs from the ISFinder database (2024-06-18) [48, 49]. Genes with ≥80% sequence identity and ≥20 amino acid overlap to any IS were discarded, resulting in the exclusion of 18 ARG sequences. Next, all ARG sequences were clustered at 90% nucleotide similarity using VSEARCH v2.28.1 [50] to remove redundancy. We then used BLASTp v2.15.0 [47] to align the proteins encoded by the resulting centroid sequences against sequences from the ResFinder database of acquired ARGs (downloaded 2024-06-19) [6]. Genes displaying ≥90% sequence identity and ≥70% coverage to any ResFinder sequence were classified as “Established”, otherwise they were classified as “Latent”. In total, this resulted in a reference database of 720 established ARGs and 33,224 latent ARGs.

### Retrieval of metagenomic data

Human gut metagenomic samples were downloaded from the MGnify database (2024-07-23) [51]. Here, only data generated using Illumina platforms were included to ensure comparability. Samples were excluded if the study had fewer than five sequencing runs in total or if unique links to fastq files in the European Nucleotide Archive API (ENA) [52] were missing. Finally, samples representing the gut microbiome of infant humans were removed due to the differences between infant and adult gut microbiome composition. Wastewater samples were retrieved from the National Center for Biotechnology Information Sequence Read Archive (NCBI SRA) [53] based on accession numbers provided by Munk et al. 2022 [54] and Hendriksen et al. 2019 [55]. This resulted in a total of 5,694 human gut samples as well as 1,108 wastewater samples. Next, BBDuk from BBMap v39.06 [56] was used to quality control the samples, with trim quality 20, minimum length 60, and left and right trimming of the raw files. Finally, all samples containing less than 5,000,000 reads after the quality control were discarded to ensure sufficient sequencing depth, after which 5,630 human gut samples and 1,034 wastewater samples remained.

### Quantification of ARGs in metagenomes

To quantify ARGs in different environments, metagenomic forward reads were aligned to the reference ARG database using DIAMOND blastx v2.1.19 [57]. To avoid double-counting, only forward reads were considered. Reads displaying ≥95% sequence identity, ≥20% read coverage, and a minimum match length of 20 amino acids were considered matches. For reads matching multiple ARGs, the best match was selected based on 1) sequence identity, 2) alignment length, and 3) *e*-value. If all these were equal, one of the matching ARGs was randomly selected. To ensure comparability, each sample was rarefied to 5,000,000 reads. To visualize the relative abundance of specific ARGs in different environments, the rarefied counts were further normalized to log counts per million bacteria (logCPM_B_). Here, the normalized abundance *x*_*ij*_ of gene *i* in metagenomic sample *j* was calculated as

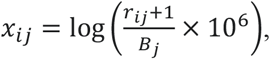

where *r*_*ij*_ is the number of reads matching gene *i*, while *B*_*j*_ is the total number of reads mapping to bacteria in sample *j*. The number of reads mapping to bacteria was calculated by applying Kraken2 v2.1.3 to each of the quality-controlled metagenomic samples, with parameters --paired –confidence 0.1 --use-names [58].

Next, we estimated the prevalence of each ARG in the human gut and wastewater by calculating the proportion of samples from each environment where the ARG was present, here defined as ≥3 matching reads. Based on these results, the ARGs were divided into four categories. Here, ARGs that were present in ≥5% of both human gut and wastewater samples were classified as co-promoted, while genes that were present in ≥5% of samples from only one type of environment were classified as human gut (HG)-promoted or wastewater (WW)-promoted, respectively. Finally, ARGs that were not present in any metagenomic sample were removed from consideration and the remaining genes that were present in <5% of both human and wastewater samples were classified as non-promoted.

### Taxonomic analysis of bacterial hosts

For each ARG, the taxonomy of the bacterial genome(s) in which it was identified was recorded at each taxonomic level. To further increase the reliability of the results, we required at least 3 observations of a taxon to record it for ARGs that were identified in ≥3 genomes; for ARGs found in fewer genomes, the required number of taxon observations was the same as the identified number of host genomes. Based on the observed host taxonomies, for each ARG, we identified the lowest taxonomic level at which all known host species of the ARG shared a common ancestor, and used this to estimate the host range of established and latent ARGs across the four promotion categories. To enable comparisons between categories, each ARG category was first subsampled to match the size of the smallest category, after which we calculated the proportion of subsampled ARGs whose most recent common ancestor was found at each taxonomic level. This procedure was repeated 1,000 times and the mean over the observed proportions was calculated. To not remove too much data during the subsampling step, categories with <10 genes were excluded from the analysis. To assess overrepresentation of specific ARGs among genes with a given host-range, we performed a permutation test by randomly shuffling the category labels of each ARG, and repeated the procedure described above. This was then repeated 1,000 times, after which *p*-values were calculated as the proportion of generated values that were at least as extreme as the true observation.

To assess the presence of ARGs in pathogenic bacteria, the identified genomic host species were cross-referenced against a list of 15 species associated with high mortality rates in clinical infections compiled based on Ikuta et al. 2022 [18]. We then analyzed whether established and latent genes from each category were overrepresented among ARGs carried by ≥*n* different pathogenic host species, where *n* was any number between zero and 15 (the number of included pathogens). Here, for each *n*, the four categories were again subsampled down to the size of the smallest category and the proportion of subsampled genes from each category carried by ≥*n* different pathogens was calculated. As with the previous analysis, we excluded categories with <10 genes and extracted the mean proportions over 1,000 iterations. Here, we again assessed overrepresentation by performing 1,000 additional rounds of analysis, each time randomly permuting the category labels in the dataset. These results were then visualized and compared to the curve produced by the true observations (Supplementary Fig. 5).

### Genetic context analysis

To identify mobile genetic elements associated with environment-specific ARGs, we implemented a genetic context analysis. For each of the co-promoted, HG-promoted, WW-promoted, and non-promoted ARGs, we first retrieved genetic regions of up to 10,000 base pairs up- and downstream of the ARG from the genomes in which it was originally identified by fARGene using GEnView v0.2 [59]. Here, if an ARG was identified in more than 1,000 genomes, the context was only retrieved from 1,000 randomly selected genomes to improve computational efficiency. The retrieved contexts were then screened for genes associated with mobile genetic elements as well as co-localized mobile ARGs. To identify genes involved in plasmid conjugation, the genetic regions were translated in all six reading frames using EMBOSS Transeq v6.5.7 [60] and screened with 124 hidden Markov models from MacSyfinder Conjscan v2.0 [61] using HMMER v3.1b2 [62]. Next, co-localized mobile ARGs were identified by using BLASTn v2.10.1 [63] to align the genetic regions against the ResFinder database [6], and finding the best (highest sequence identity) among overlapping hits with >90% nucleotide identity and >75% coverage to a known ARG.

### Analysis of genetic compatibility

Next, we estimated the genetic compatibility between ARGs from the identified categories and the included pathogens by calculating the nucleotide composition dissimilarity. This was computed by first selecting a representative genome for each pathogen from NCBI RefSeq [64]. The nucleotide sequences of the ARGs and genomes were divided into 5-mers, and their respective 5-mer distributions were computed. Then, we calculated the Euclidean distances between the 5-mer distributions of the ARGs in each category and each representative pathogen genome in turn. For each promotion category and pathogen, the resulting distributions of gene-genome 5-mer distances were visualized as boxplots, and the median distances were extracted and visualized together with the proportion of ARGs from that category carried by that pathogen.

We then expanded this analysis to estimate the genetic compatibility between differently promoted ARGs and bacteria that are specific to the human gut or wastewater. The human gut- and wastewater-specific bacterial genera were identified based on analysis of each quality-controlled metagenomic sample with Kraken2 v2.1.3, with parameters --paired --confidence 0.1 --use-names [58]. A bacterial genus was considered present in a sample if >0.01% of the reads mapped to that genus. Then, we identified genera which were present in ≥5% of human gut samples and ≥5% of wastewater samples, which were classified as universal; genera which were present in ≥5% of human gut samples and <5% of wastewater samples, which were classified as human gut-specific; and genera which were present in <5% of human gut samples and ≥5% of wastewater samples, which were classified as wastewater-specific. For each of the included genera, we then randomly selected up to ten individual species and retrieved their representative genomes from NCBI RefSeq [64]. Genera for which <5 species were available were discarded. Finally, the genetic compatibility between each of these genomes and the ARGs from each promotion category was computed as described above.

## Data availability

All raw data analyzed in this study have been retrieved from the public repositories NCBI [40, 53] and MGnify [51]. Accession numbers of the analyzed genomes and metagenomes are provided in Supplementary Data 1.

## Code availability

Scripts used to perform the analyses and generate all figures presented in this paper are available via GitHub at https://github.com/davidgllund/promoted_args_hg_ww.

## Acknowledgements

This research was supported by the Swedish Research Council (VR) (2018–02835, 2018-05771,2019– 03482, and 2022-00945). J.B-P. acknowledges funding from the Swedish Research Council (VR; grant 2023-01721) under the frame of JPI AMR (SEARCHER; JPIAMR2023-DISTOMOS-016), the Data-Driven Life Science (DDLS) program supported by the Knut and Alice Wallenberg Foundation (KAW 2020.0239), and the Swedish Foundation for Strategic Research (FFL21-0174). Funding sources took no part in the design, analysis, or interpretation of the results.

## Author contributions

D.L., A.J., M.H., J.S.I-D., J.B-P., and E.K. designed the study and developed the approach. D.L. downloaded the bacterial genomes and generated the reference ARG database. D.L. and M.H. retrieved the metagenomic data and analyzed it for ARG abundance. D.L. performed all additional computational analysis. All authors. discussed the results and their implications. D.L. and E.K. drafted and edited the manuscript. All authors edited and approved the final manuscript.

## References

1. Murray, C.J., et al., Global burden of bacterial antimicrobial resistance in 2019: a systematic analysis. The lancet, 2022. 399(10325): p. 629–655.

2. Brito, I.L., Examining horizontal gene transfer in microbial communities. Nature Reviews Microbiology, 2021. 19(7): p. 442–453.

3. Parras-Moltó, M., et al., The transfer of antibiotic resistance genes between evolutionary distant bacteria. bioRxiv, 2024: p. 2024.10.22.619579.

4. Groussin, M., et al., Elevated rates of horizontal gene transfer in the industrialized human microbiome. Cell, 2021. 184(8): p. 2053–2067. e18.

5. Alcock, B.P., et al., CARD 2023: expanded curation, support for machine learning, and resistome prediction at the Comprehensive Antibiotic Resistance Database. Nucleic acids research, 2023. 51(D1): p. D690–D699.

6. Bortolaia, V., et al., ResFinder 4.0 for predictions of phenotypes from genotypes. Journal of Antimicrobial Chemotherapy, 2020. 75(12): p. 3491–3500.

7. Inda-Díaz, J.S., et al., Latent antibiotic resistance genes are abundant, diverse, and mobile in human, animal, and environmental microbiomes. Microbiome, 2023. 11(1): p. 44.

8. Yin, X., et al., Global environmental resistome: distinction and connectivity across diverse habitats benchmarked by metagenomic analyses. Water research, 2023. 235: p. 119875.

9. Nesme, J., et al., Large-scale metagenomic-based study of antibiotic resistance in the environment. Current biology, 2014. 24(10): p. 1096–1100.

10. Pal, C., Bengtsson-Palme, J., Kristiansson, E., and Larsson, D.G.J., The structure and diversity of human, animal and environmental resistomes. Microbiome, 2016. 4: p. 1–15.

11. Che, Y., et al., Mobile antibiotic resistome in wastewater treatment plants revealed by Nanopore metagenomic sequencing. Microbiome, 2019. 7: p. 1–13.

12. Karkman, A., Berglund, F., Flach, C.-F., Kristiansson, E., and Larsson, D.J., Predicting clinical resistance prevalence using sewage metagenomic data. Communications Biology, 2020. 3(1): p. 711.

13. Foxman, B., et al., Wastewater surveillance of antibiotic-resistant bacteria for public health action: potential and challenges. American Journal of Epidemiology, 2025. 194(5): p. 1192–1199.

14. Zhang, Z., et al., Assessment of global health risk of antibiotic resistance genes. Nature communications, 2022. 13(1): p. 1553.

15. Zhang, A.-N., et al., An omics-based framework for assessing the health risk of antimicrobial resistance genes. Nature communications, 2021. 12(1): p. 4765.

16. Garner, E., et al., Towards risk assessment for antibiotic resistant pathogens in recycled water: a systematic review and summary of research needs. Environmental microbiology, 2021. 23(12): p. 7355–7372.

17. Calderón-Franco, D., et al., Metagenomic profiling and transfer dynamics of antibiotic resistance determinants in a full-scale granular sludge wastewater treatment plant. Water Research, 2022. 219: p. 118571.

18. Ikuta, K.S., et al., Global mortality associated with 33 bacterial pathogens in 2019: a systematic analysis for the Global Burden of Disease Study 2019. The Lancet, 2022. 400(10369): p. 2221–2248.

19. Lund, D., et al., Genetic compatibility and ecological connectivity drive the dissemination of antibiotic resistance genes. Nature Communications, 2025. 16(1): p. 2595.

20. Baquero, F., et al., Evolutionary pathways and trajectories in antibiotic resistance. Clinical Microbiology Reviews, 2021. 34(4): p. e00050–19.

21. Coluzzi, C., Garcillán-Barcia, M.P., de la Cruz, F., and Rocha, E.P., Evolution of plasmid mobility: origin and fate of conjugative and nonconjugative plasmids. Molecular Biology and Evolution, 2022. 39(6): p. msac115.

22. Cury, J., Touchon, M., and Rocha, E.P., Integrative and conjugative elements and their hosts: composition, distribution and organization. Nucleic acids research, 2017. 45(15): p. 8943–8956.

23. Smillie, C., Garcillán-Barcia, M.P., Francia, M.V., Rocha, E.P., and de la Cruz, F., Mobility of plasmids. Microbiology and Molecular Biology Reviews, 2010. 74(3): p. 434–452.

24. Gingold, H. and Pilpel, Y., Determinants of translation efficiency and accuracy. Molecular systems biology, 2011. 7(1): p. 481.

25. Hughes, D. and Andersson, D.I., Environmental and genetic modulation of the phenotypic expression of antibiotic resistance. FEMS microbiology reviews, 2017. 41(3): p. 374–391.

26. Parvathy, S.T., Udayasuriyan, V., and Bhadana, V., Codon usage bias. Molecular biology reports, 2022. 49(1): p. 539–565.

27. Smillie, C.S., et al., Ecology drives a global network of gene exchange connecting the human microbiome. Nature, 2011. 480(7376): p. 241–244.

28. Crits-Christoph, A., Hallowell, H.A., Koutouvalis, K., and Suez, J., Good microbes, bad genes? The dissemination of antimicrobial resistance in the human microbiome. Gut microbes, 2022. 14(1): p. 2055944.

29. Forster, S.C., et al., Strain-level characterization of broad host range mobile genetic elements transferring antibiotic resistance from the human microbiome. Nature Communications, 2022. 13(1): p. 1445.

30. Fang, G.-Y., Liu, X.-Q., Jiang, Y.-J., Mu, X.-J., and Huang, B.-W., Horizontal gene transfer in activated sludge enhances microbial antimicrobial resistance and virulence. Science of The Total Environment, 2024. 912: p. 168908.

31. Ebmeyer, S., Kristiansson, E., and Larsson, D.J., A framework for identifying the recent origins of mobile antibiotic resistance genes. Communications biology, 2021. 4(1): p. 8.

32. Ebmeyer, S., Kristiansson, E., and Larsson, D.J., Unraveling the origins of mobile antibiotic resistance genes using random forest classification of large-scale genomic data. Environment International, 2025: p. 109374.

33. Philippon, A., Labia, R., and Jacoby, G., Extended-spectrum beta-lactamases. Antimicrobial agents and chemotherapy, 1989. 33(8): p. 1131–1136.

34. Speer, B.S., Shoemaker, N.B., and Salyers, A.A., Bacterial resistance to tetracycline: mechanisms, transfer, and clinical significance. Clinical microbiology reviews, 1992. 5(4): p. 387–399.

35. Roberts, M.C., et al., Nomenclature for macrolide and macrolide-lincosamide-streptogramin B resistance determinants. Antimicrobial agents and chemotherapy, 1999. 43(12): p. 2823–2830.

36. McInnes, R.S., McCallum, G.E., Lamberte, L.E., and van Schaik, W., Horizontal transfer of antibiotic resistance genes in the human gut microbiome. Current opinion in microbiology, 2020. 53: p. 35–43.

37. Karkman, A., Do, T.T., Walsh, F., and Virta, M.P., Antibiotic-resistance genes in waste water. Trends in microbiology, 2018. 26(3): p. 220–228.

38. Brenciani, A., et al., Genetic elements carrying erm (B) in Streptococcus pyogenes and association with tet (M) tetracycline resistance gene. Antimicrobial agents and chemotherapy, 2007. 51(4): p. 1209–1216.

39. Crofts, T.S., Gasparrini, A.J., and Dantas, G., Next-generation approaches to understand and combat the antibiotic resistome. Nature Reviews Microbiology, 2017. 15(7): p. 422–434.

40. Kitts, P.A., et al., Assembly: a resource for assembled genomes at NCBI. Nucleic acids research, 2016. 44(D1): p. D73–D80.

41. Berglund, F., et al., Identification and reconstruction of novel antibiotic resistance genes from metagenomes. Microbiome, 2019. 7: p. 1–14.

42. Lund, D., et al., Extensive screening reveals previously undiscovered aminoglycoside resistance genes in human pathogens. Communications Biology, 2023. 6(1): p. 812.

43. Lund, D., et al., Large-scale characterization of the macrolide resistome reveals high diversity and several new pathogen-associated genes. Microbial Genomics, 2022. 8(1): p. 000770.

44. Berglund, F., et al., Comprehensive screening of genomic and metagenomic data reveals a large diversity of tetracycline resistance genes. Microbial genomics, 2020. 6(11): p. e000455.

45. Berglund, F., et al., Identification of 76 novel B1 metallo-β-lactamases through large-scale screening of genomic and metagenomic data. Microbiome, 2017. 5: p. 1–13.

46. Boulund, F., Johnning, A., Pereira, M.B., Larsson, D.G.J., and Kristiansson, E., A novel method to discover fluoroquinolone antibiotic resistance (qnr) genes in fragmented nucleotide sequences. BMC genomics, 2012. 13: p. 1–9.

47. Camacho, C., et al., BLAST+: architecture and applications. BMC bioinformatics, 2009. 10: p. 1–9.

48. Siguier, P., Pérochon, J., Lestrade, L., Mahillon, J., and Chandler, M., ISfinder: the reference centre for bacterial insertion sequences. Nucleic acids research, 2006. 34(suppl_1): p. D32–D36.

49. Insertion Sequence (IS) database. 2024-06-18]; Available from: https://github.com/thanhleviet/ISfinder-sequences.

50. Rognes, T., Flouri, T., Nichols, B., Quince, C., and Mahé, F., VSEARCH: a versatile open source tool for metagenomics. PeerJ, 2016. 4: p. e2584.

51. Richardson, L., et al., MGnify: the microbiome sequence data analysis resource in 2023. Nucleic Acids Research, 2023. 51(D1): p. D753–D759.

52. Yuan, D., et al., The European nucleotide archive in 2023. Nucleic Acids Research, 2024. 52(D1): p. D92–D97.

53. Leinonen, R., Sugawara, H., Shumway, M., and Collaboration, I.N.S.D., The sequence read archive. Nucleic acids research, 2010. 39(suppl_1): p. D19–D21.

54. Munk, P., et al., Genomic analysis of sewage from 101 countries reveals global landscape of antimicrobial resistance. Nature Communications, 2022. 13(1): p. 7251.

55. Hendriksen, R.S., et al., Global monitoring of antimicrobial resistance based on metagenomics analyses of urban sewage. Nature communications, 2019. 10(1): p. 1124.

56. Bushnell, B., BBMap: a fast, accurate, splice-aware aligner. 2014.

57. Buchfink, B., Xie, C., and Huson, D.H., Fast and sensitive protein alignment using DIAMOND. Nature methods, 2015. 12(1): p. 59–60.

58. Wood, D.E., Lu, J., and Langmead, B., Improved metagenomic analysis with Kraken 2. Genome biology, 2019. 20: p. 1–13.

59. Ebmeyer, S., Coertze, R.D., Berglund, F., Kristiansson, E., and Larsson, D.G.J., GEnView: a gene-centric, phylogeny-based comparative genomics pipeline for bacterial genomes and plasmids. Bioinformatics, 2022. 38(6): p. 1727–1728.

60. Madeira, F., et al., The EMBL-EBI search and sequence analysis tools APIs in 2019. Nucleic acids research, 2019. 47(W1): p. W636–W641.

61. Abby, S.S., et al., Identification of protein secretion systems in bacterial genomes. Scientific reports, 2016. 6(1): p. 23080.

62. Eddy, S.R., Accelerated profile HMM searches. PLoS computational biology, 2011. 7(10): p. e1002195.

63. Altschul, S.F., Gish, W., Miller, W., Myers, E.W., and Lipman, D.J., Basic local alignment search tool. Journal of molecular biology, 1990. 215(3): p. 403–410.

64. O’Leary, N.A., et al., Reference sequence (RefSeq) database at NCBI: current status, taxonomic expansion, and functional annotation. Nucleic acids research, 2016. 44(D1): p. D733–D745.

