## Supplementary figures for "Community-promoted antibiotic resistance genes show increased dissemination among pathogens"

### Supplementary material

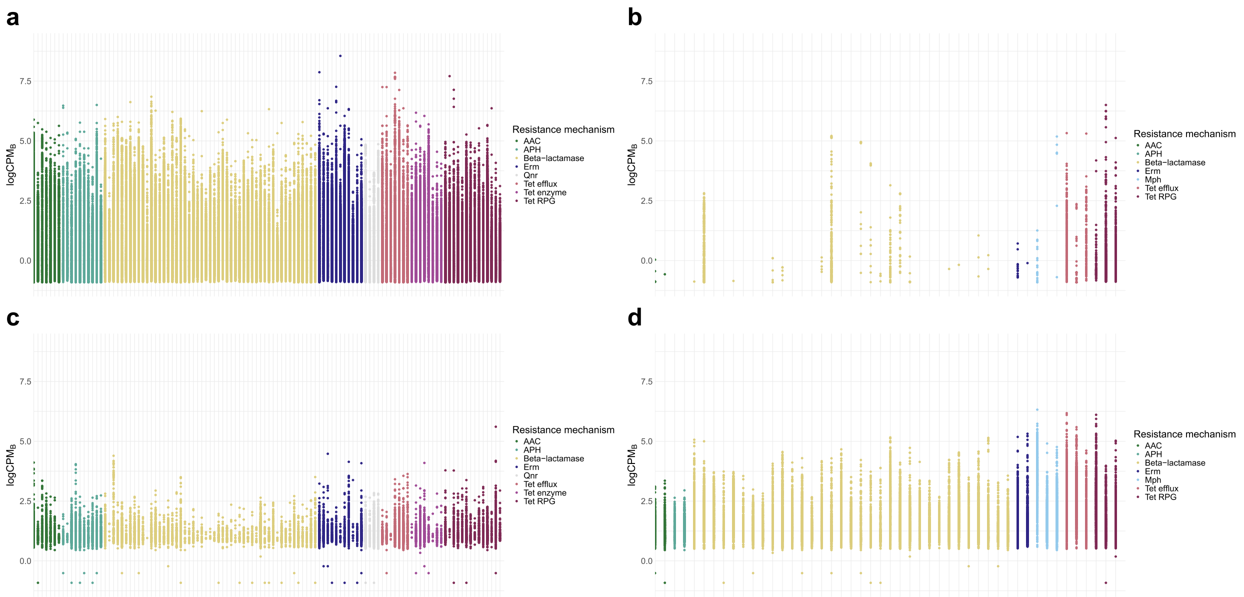


**Supplementary Fig. 1.** Abundance of human gut (HG)-promoted and wastewater (WW)-promoted antibiotic resistance genes in metagenomic samples. Each gene in each sample is rarefied and normalized by the number of reads mapping to bacteria for that sample. Instances where the gene was not detected have been removed. **a** Abundance of HG-promoted ARGs in human gut samples. **b** Abundance of WW-promoted ARGs in human gut samples. **c** Abundance of HG-promoted ARGs in wastewater samples. **d** Abundance of WW-promoted ARGs in wastewater samples.

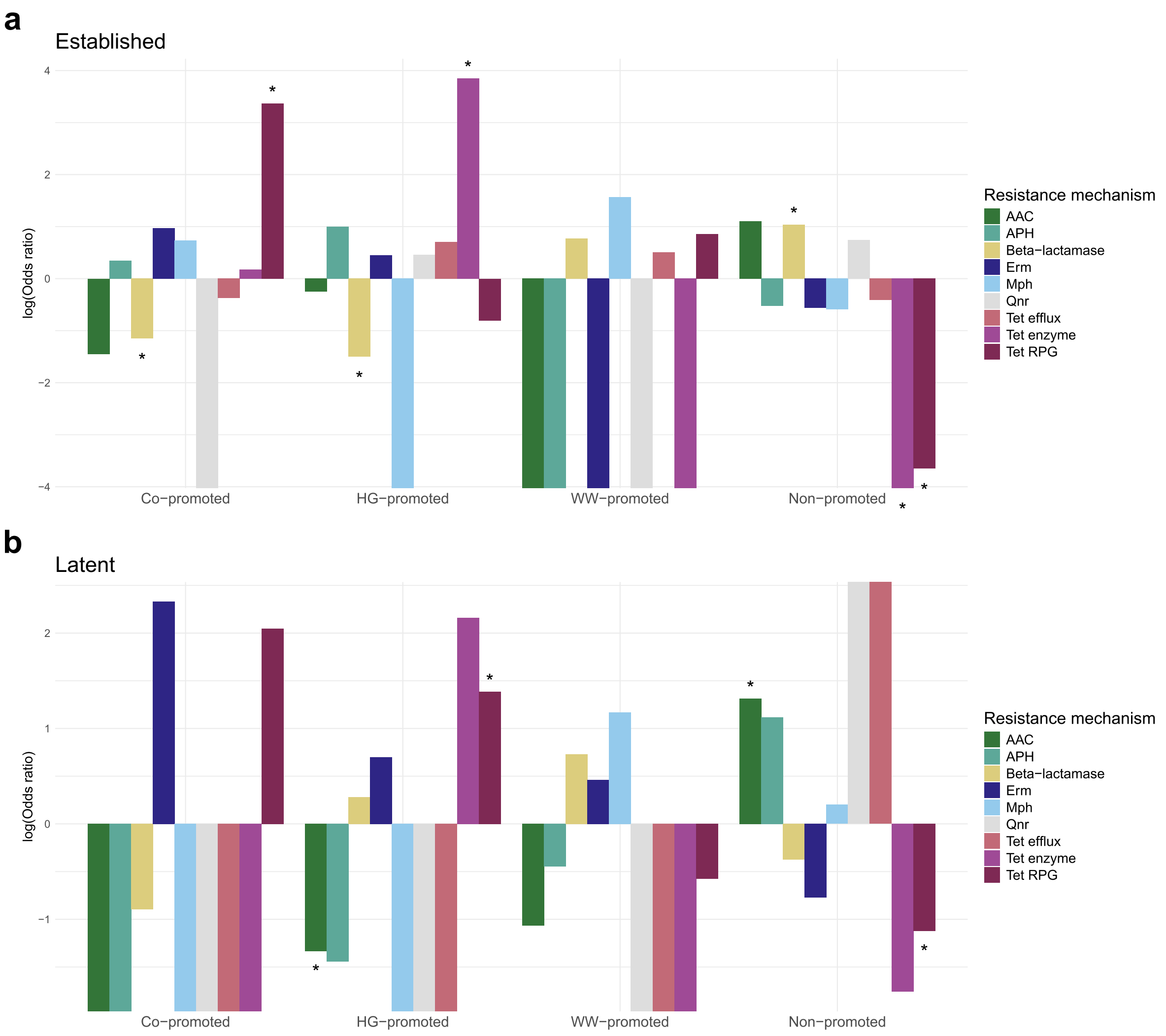


**Supplementary Fig 2.** Enrichment analysis of different resistance mechanisms among the **a** established and **b** latent ARGs from each promotion category. Odds ratios and corresponding *p*-values were calculated using Fisher's exact test. *$p<0.01$.

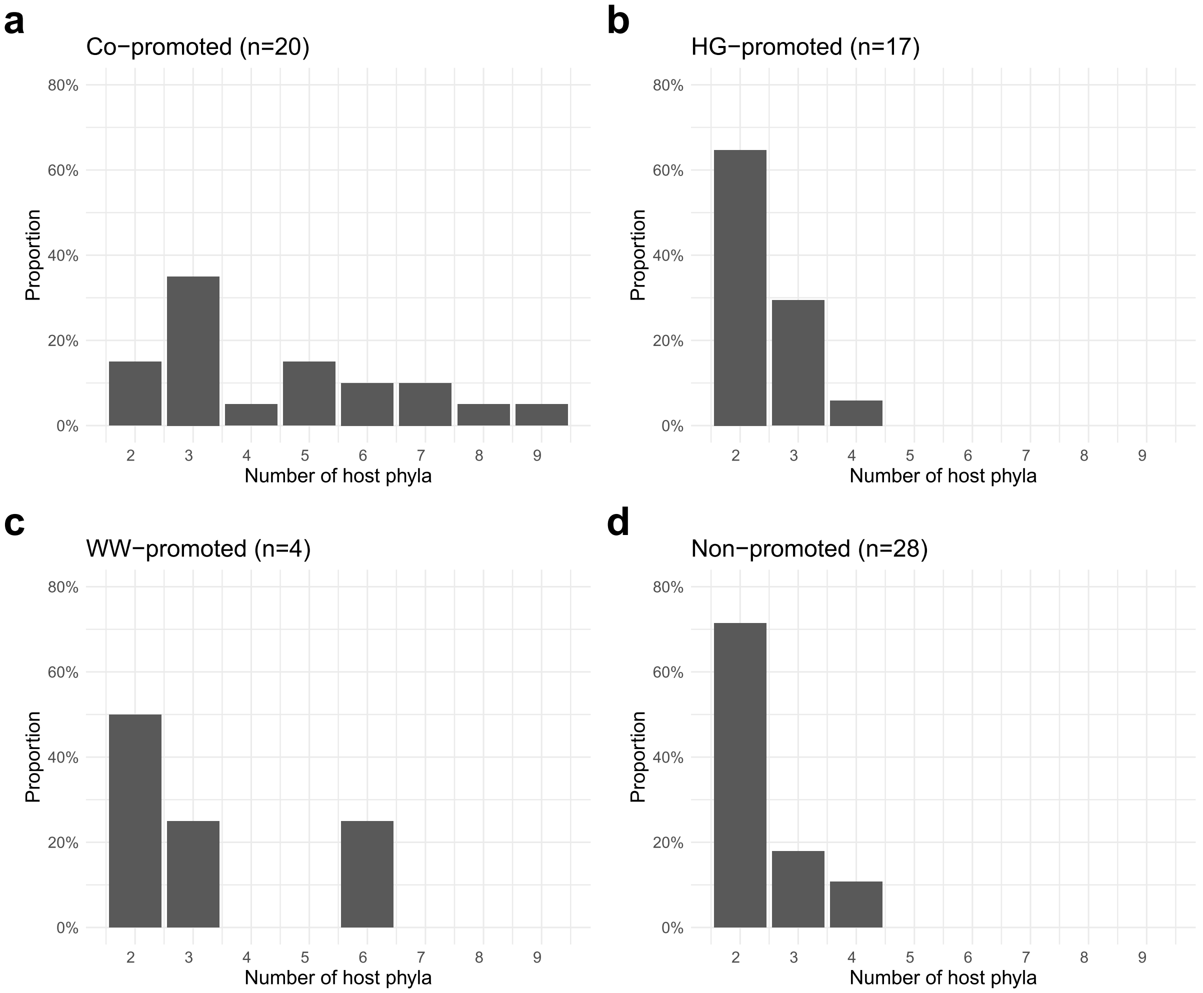


**Supplementary Fig 3.** Proportion of inter-phyla transferred ARGs from different promotion categories that were identified in any number of bacterial phyla. **a** Co-promoted ARGs. **b** HG-promoted ARGs. **c** WW-promoted ARGs. **d** Non-promoted ARGs.

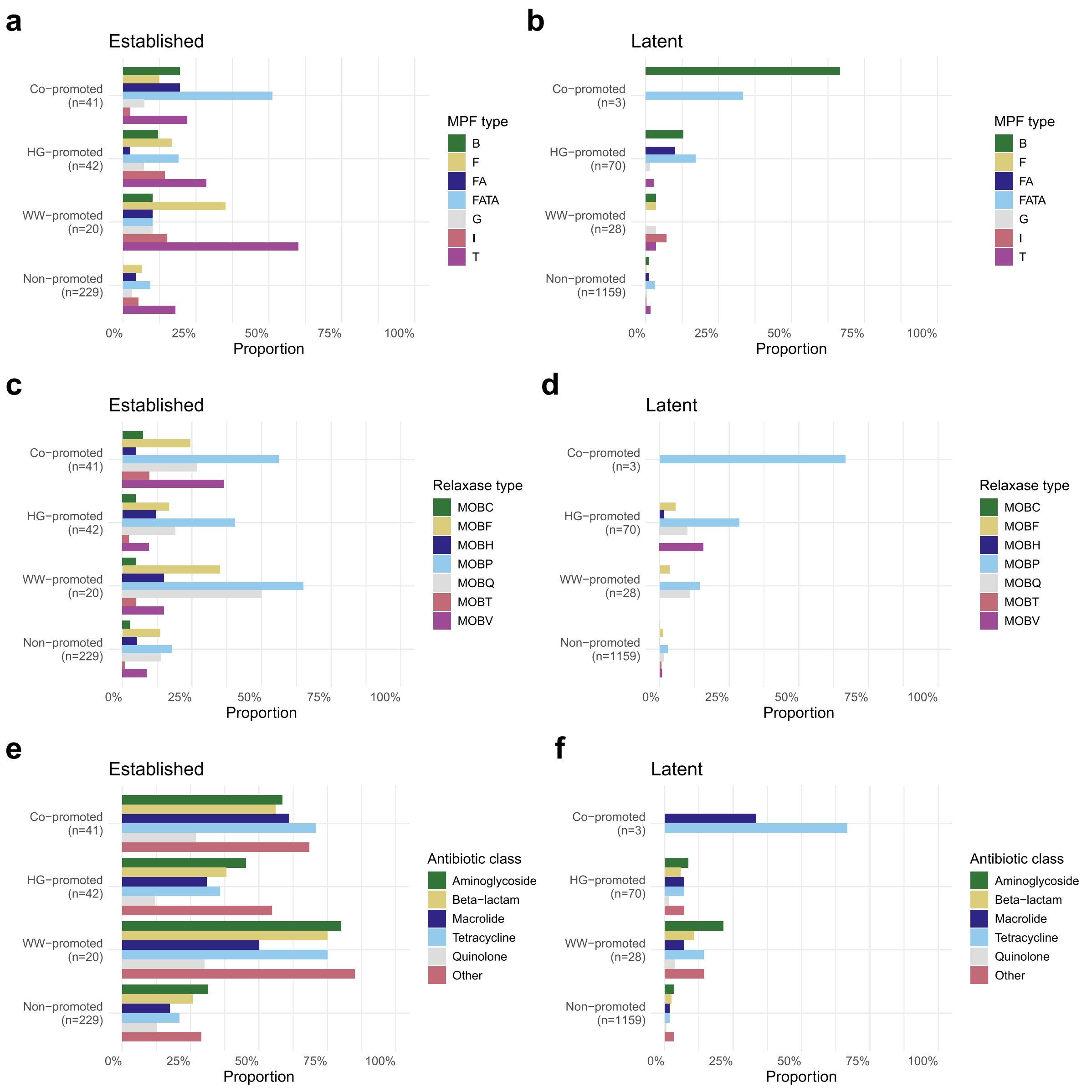


**Supplementary Fig. 4.** Distribution of mobile genetic elements (MGEs) co-localized with co-promoted, HG-promoted, WW-promoted, and non-promoted ARGs, separated into established and latent variants. For each ARG category, bar lengths indicate the proportion of ARGs with a co-localized MGE detected in at least one host genome. Panels (**a**-**b**) show the distribution of mating pair formation (MPF) genes located near established and latent ARGs across categories. Panels (**c**-**d**) display relaxase genes, and panels (**e**-**f**) show other mobile ARGs, all co-localized with established and latent ARGs from the different categories.

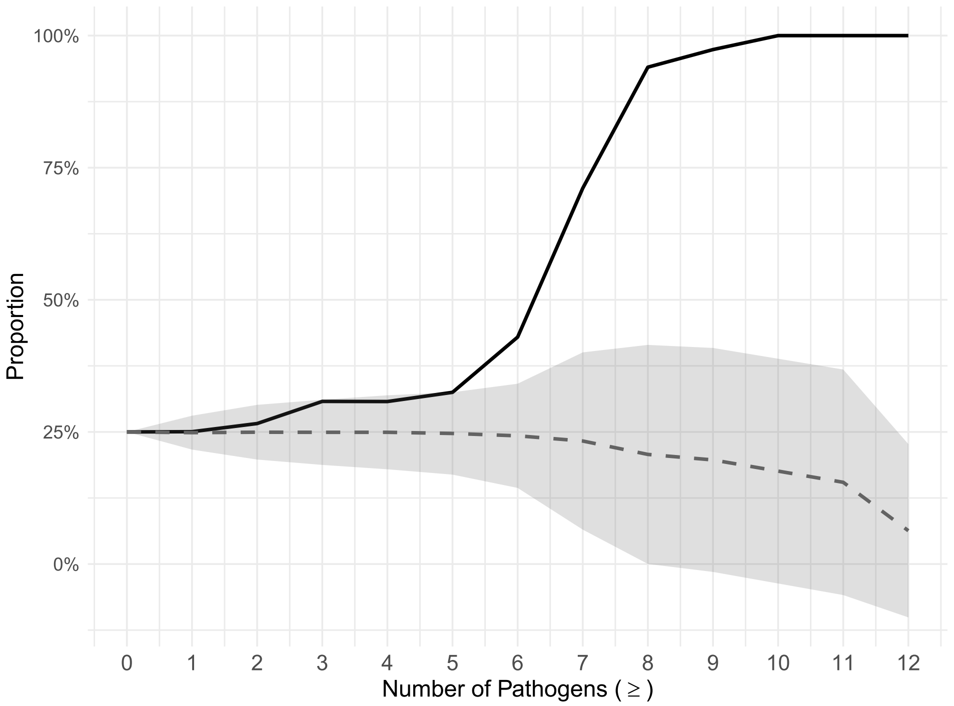


**Supplementary Fig. 5.** Overrepresentation of co-promoted genes among ARGs identified in multiple pathogens. The solid line depicts the mean proportion of established ARGs carried by ≥n pathogens made up of co-promoted genes after 1,000 repetitions of subsampling the four promotion categories to equal size (for details, see Methods). The dashed line and ribbon represent the mean ± SD produced by repeating the analysis 1,000 times, each time permuting the labels denoting the genes' promotion category (for details, see Methods).

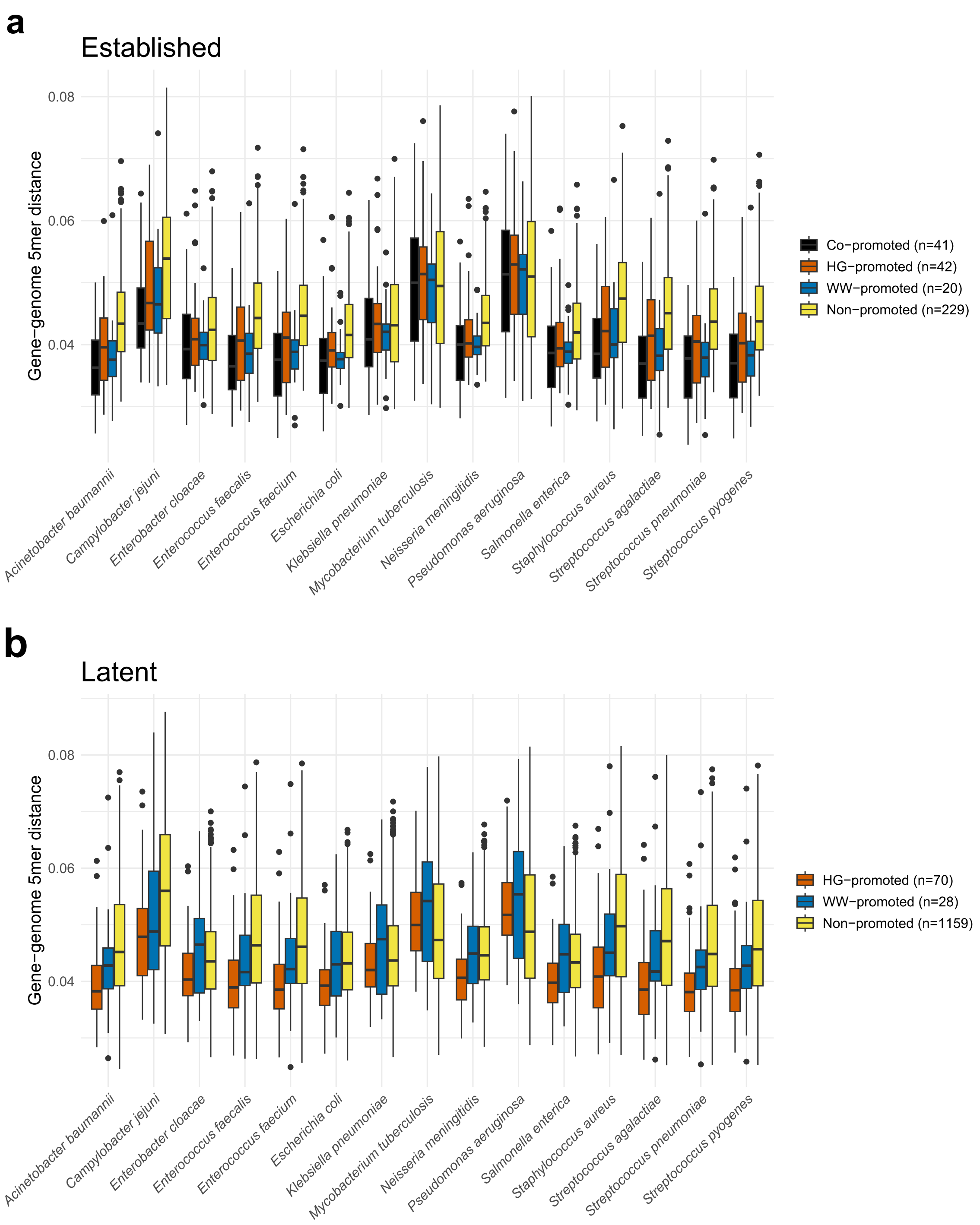


**Supplementary Fig. 6.** Euclidean distances between the 5mer distributions of **a** established, and **b** latent ARGs and representative genomes from bacterial pathogens. The ARGs have been divided into the four categories (co-promoted, HG-promoted, WW-promoted, and non-promoted), and the distribution of 5mer distances between the ARGs in each category and each genome is visualized as boxplots.

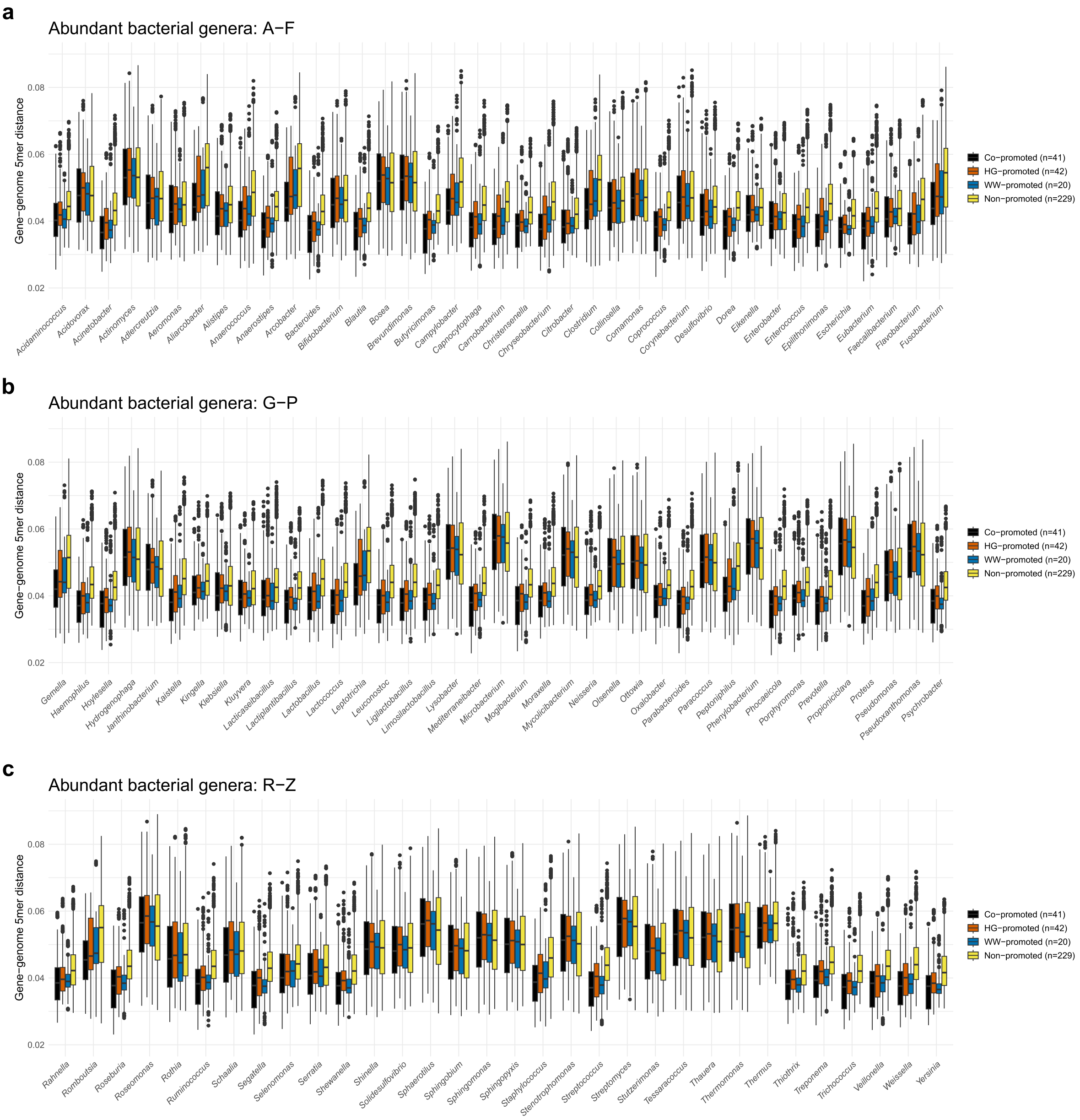


**Supplementary Fig. 7.** Euclidean distances between the 5mer distributions of established ARGs and representative genomes from bacterial genera that were abundant in human gut and wastewater metagenomic samples. The ARGs have been divided into the four categories (co-promoted, HG-promoted, WW-promoted, and non-promoted), and the distribution of 5mer distances between the ARGs in each category and each genome is visualized as boxplots. Each bacterial genus is represented by up to ten representative genomes, each from a unique species.

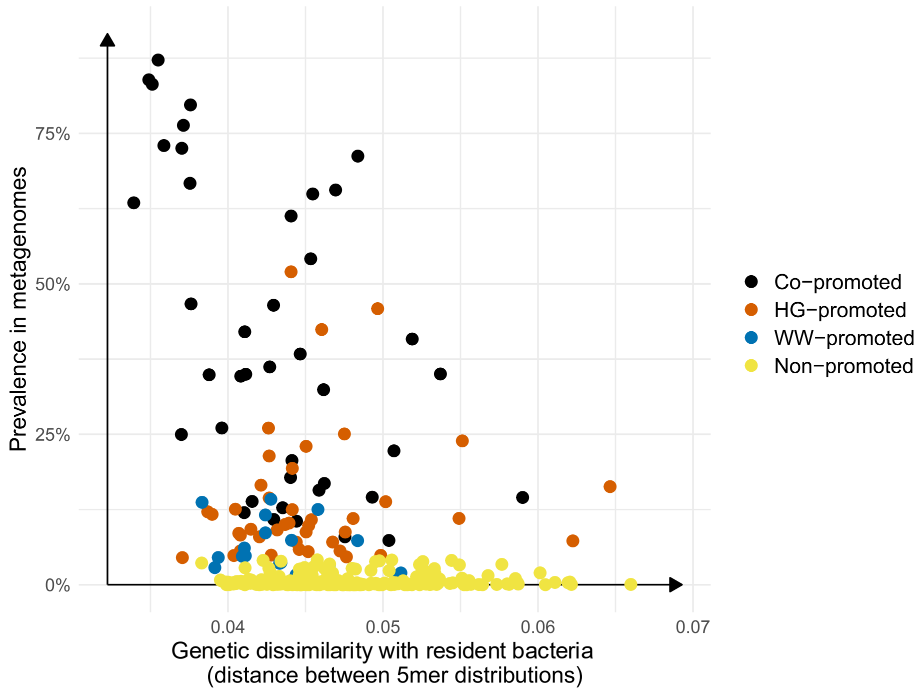


**Supplementary Fig. 8.** Relationship between genetic dissimilarity with bacteria inhabiting the human gut and/or wastewater microbiome(s) and prevalence in these environments. Each point represents an established ARG, with the *y*-axis representing its prevalence among the analyzed metagenomic samples, and the x-axis representing the mean of its median genetic incompatibilities with a total of 110 bacterial genera frequently encountered in the human gut and/or wastewater. The colors denote the promotion category of each ARG.

**Supplementary Data 1.** Accession IDs and Sample IDs of the genomes and metagenomes analyzed in this study.

*Available with the online version of the paper*
